# Increased mitochondrial transcription initiation does not promote oxidative phosphorylation

**DOI:** 10.1101/2024.01.11.575236

**Authors:** Maria Miranda, Andrea Mesaros, Irina Kuznetsova, Martin Purrio, Louise Pérard, Aleksandra Filipovska, Arnaud Mourier, Nils-Göran Larsson, Inge Kühl

## Abstract

POLRMT is the sole RNA polymerase in human mitochondria where it generates primers for mitochondrial DNA (mtDNA) replication and transcribes the mitochondrial genome to express genes encoding essential components of the oxidative phosphorylation (OXPHOS) system. Elevated POLRMT levels are found in several cancers and in mouse models with severe mitochondrial dysfunction. Here, we generated and characterized mice over-expressing *Polrmt* to investigate the physiological and molecular consequences of elevated POLRMT levels. Increasing POLRMT did not result in any pathological phenotype but instead positively affected exercise capacity under stress conditions. POLRMT overexpression increased *in organello* transcription initiation, resulting in higher steady-state levels of the promoter-proximal L-strand transcript 7S RNA and higher mtDNA levels. Surprisingly, the abundance of mature mitochondrial RNAs was not affected by the elevated POLRMT levels. Furthermore, ubiquitous simultaneous overexpression of POLRMT and LRPPRC, which stabilizes mitochondrial messenger RNAs, did not increase steady-state levels of mitochondrial transcripts in the mouse. Our data show that POLRMT levels regulate transcription initiation, but additional regulatory steps downstream of transcription initiation and transcript stability limit OXPHOS biogenesis.

## INTRODUCTION

Biogenesis of the mitochondrial oxidative phosphorylation (OXPHOS) system is vital for mammalian life as it fulfils several metabolic functions, including the generation of most cellular energy in the form of ATP, and OXPHOS is altered in a broad range of human pathologies, including cancer and aging. The OXPHOS system is under the dual genetic control of the nuclear and mitochondrial genomes (nDNA and mtDNA, respectively)(Gustafsson et al., 2016). In mammals, the nDNA encodes most mitochondrial proteins, including all factors required to express and maintain mtDNA and ∼90 OXPHOS subunits, while the mtDNA encodes 13 essential subunits of four out of the five OXPHOS complexes (Rath et al., 2021), two mitochondrial ribosomal RNAs (mt-rRNAs) and 22 transfer RNAs (mt-tRNAs). Failure to express mtDNA causes a global loss of mitochondrial protein complexes with dual genetic contributions, i.e., the OXPHOS complexes and the mitochondrial ribosome (Kühl et al., 2017), suggesting that regulation of mtDNA gene expression is not only fundamental for proper mitochondrial function but also allows local and rapid adaptations to bioenergetic and metabolic demands.

Mammalian mtDNA is an intron-free, circular molecule present in multiple copies per cell and it is composed of two coding strands known as the heavy (H) and the light (L) strand because of differences in their G+T base composition(Gustafsson et al., 2016). Additionally, mammalian mtDNA has a non-coding control region of ∼1kb (NCR) which contains the H-strand origin of replication (OH) and at least two promoters for transcription of the H and L strands (HSP and LSP, respectively)(Chang et al., 1985; Crews et al., 1979; Tan et al., 2022). Transcription of mtDNA generates near genome-length polycistronic transcripts that are processed to yield individual, mature RNA molecules (mt-RNAs)(Holzmann et al., 2008; Montoya et al., 1982; Ojala et al., 1981; Rackham et al., 2016; Siira et al., 2018), and stabilized by RNA-binding proteins. The link between post-transcriptional RNA processing and translation is not fully known, but it has been shown that the leucine-rich pentatricopeptide repeat-containing protein (LRPPRC) promotes polyadenylation and stabilizes all mt-mRNAs except *mt-Nd6* (Gohil et al., 2010; Harmel et al., 2013; Ruzzenente et al., 2012; Sasarman et al., 2010).

Although the basic machinery of mtDNA transcription has been thoroughly studied *in vitro* (Hillen et al., 2018), little is known about its regulation *in vivo*. Transcription of mammalian mtDNA is performed by a nuclear-encoded, single-subunit mitochondrial RNA polymerase (POLRMT) that operates exclusively in mitochondria (Gustafsson et al., 2016; Kühl et al., 2016, 2014; Ringel et al., 2011). Several factors accompany POLRMT throughout the different steps of mitochondrial transcription: the mitochondrial transcription factors A and B2 (TFAM and TFB2M) that aid in promoter recognition and activation (Hillen et al., 2017), the mitochondrial transcription elongation factor (TEFM) that increases the processivity of POLRMT (Jiang et al., 2019; Minczuk et al., 2011), and the mitochondrial transcription termination factor 1 (MTERF1) that acts as a roadblock terminating L-strand transcription(Terzioglu et al., 2013). Remarkably, POLRMT also generates the primers required for mtDNA replication initiation at OL and OH (Falkenberg, 2018; Kühl et al., 2016; Sarfallah et al., 2021), placing this enzyme at the core of both mtDNA expression and maintenance. A proportion of LSP transcripts is terminated in the NCR (Wanrooij et al., 2012) to generate RNA primers for DNA synthesis at OH (Gustafsson et al., 2016). The regulation of transcription for primer formation versus gene expression is partly understood, and different factors such as mitochondrial RNase H1 and mitochondrial single-strand binding protein (SSBP1) are of key importance in this process (Agaronyan et al., 2015; Jiang et al., 2021; Kühl et al., 2016).

Recent studies have highlighted the medical importance of understanding the regulation of POLRMT and mitochondrial transcription. Pathogenic mutations in *POLRMT* cause mitochondrial dysfunction with a broad spectrum of neurodevelopmental presentations due to defective mitochondrial transcription (Oláhová et al., 2021). Elevated POLRMT levels were found in patients with lung and breast cancer (Sotgia et al., 2012) as well as in acute myeloid leukemia cells (Bralha et al., 2015; Chaudhary et al., 2021); therefore, POLRMT has been suggested to be a novel important oncogene (Miranda et al., 2022; Zhou et al., 2021). Consistently, POLRMT has been proposed as a therapeutic target to treat some cancers (Bonekamp et al., 2020; Bralha et al., 2015). Furthermore, increased POLRMT levels are found in many different mouse models with mitochondrial dysfunction (Jiang et al., 2019; Kühl et al., 2017; Milenkovic et al., 2013; Perks et al., 2018; Silva Ramos et al., 2019). To date, it remains unknown whether POLRMT contributes to pathogenesis or can increase OXPHOS under physiologic conditions. Here, we investigated the physiological and molecular consequences of elevated POLRMT levels in non-pathogenic conditions by generating and characterizing mice ubiquitously over-expressing POLRMT. The POLRMT overexpressing mice are viable and have no evident pathologic phenotype up until one year of age, the latest timepoint studied. Remarkably, overexpression of POLRMT had a positive effect on exercise capacity under exercise challenge conditions. Molecular analysis of various tissues of the POLRMT overexpressing mice suggests that POLRMT is the limiting factor for mtDNA transcription initiation because elevated POLRMT levels increase *in organello* transcription initiation and steady-state levels of the promoter-proximal 7S RNA transcript of the L-strand transcription, as well as mtDNA levels. Under normal conditions, an increase in mtDNA gene expression is not translated into an increase in OXPHOS capacity because mature mitochondrial transcripts are not globally upregulated. Our findings support that regulatory checkpoints occur downstream of transcription initiation, balancing mtDNA transcription elongation and post-transcriptional processes depending on energetic needs *in vivo*.

## RESULTS

### Overexpression of *Polrmt* results in healthy mice with normal OXPHOS

To study the physiological role of increased POLRMT levels, we generated mice ubiquitously overexpressing *Polrmt* using a bacterial artificial chromosome (BAC) transgenic strategy (Park et al., 2007). A BAC clone containing a fragment of chromosome 10 with the *Polrmt* gene was modified to introduce a silent point mutation (c420G>T) that generates a HindIII restriction site in exon 3, differentiating the transgene from the endogenous *Polrmt* alleles (Fig 1A, B and Fig S1A). This strategy allows a moderate overexpression of *Polrmt* under the control of its endogenous promoter mimicking physiological expression levels. Germline transmission and expression of the transgene was verified by PCR and subsequent HindIII restriction digest (Fig S1B). Quantification of the transgene by pyrosequencing showed that a single copy was integrated into the mouse genome (Fig S1C), which was consistent with the observed ∼33% increase in *Polrmt* transcript levels (Fig 1C) in heart and increased POLRMT protein levels in various mouse tissues such as brown adipose tissue, heart, kidney, liver or spleen (Fig 1D). Remarkably, we never obtained founders containing more than one additional copy of the *Polrmt* gene. POLRMT overexpressors were born at Mendelian ratios (Fig S2A), and had a normal body weight gain with female POLRMT overexpressor mice showing a slight but significant increase in body weight from 150 days of age (Fig. 1E). Proportions of fat and lean mass were normal (Fig S2B, C), and POLRMT overexpressor mice showed no abnormal behavior up until 52 weeks of age. Furthermore, there was no evidence of cardiomyopathy as the heart-to-body weight ratios were normal at 13 and 26 weeks (Fig S2D). No mice were euthanized due to abnormal cancerous masses during the course of the study. We next proceeded to evaluate whether moderately increased levels of POLRMT influenced the mitochondrial bioenergetic capacity. We found normal respiration of isolated mitochondria from heart and liver incubated with complex I or complex II substrates and analyzed under phosphorylating (state 3), non-phosphorylating (state 4) or uncoupled conditions (Fig 1F, G), indicating that the OXPHOS system is fully functional. To further verify our findings, we performed enzyme activity and western blot analysis for respiratory chain complexes and found no differences between wild-type and POLRMT overexpressor mice (Fig. S3A, B). These results show that a moderate increase in POLRMT levels does not affect OXPHOS function or cause pathology.

**Figure 1:**
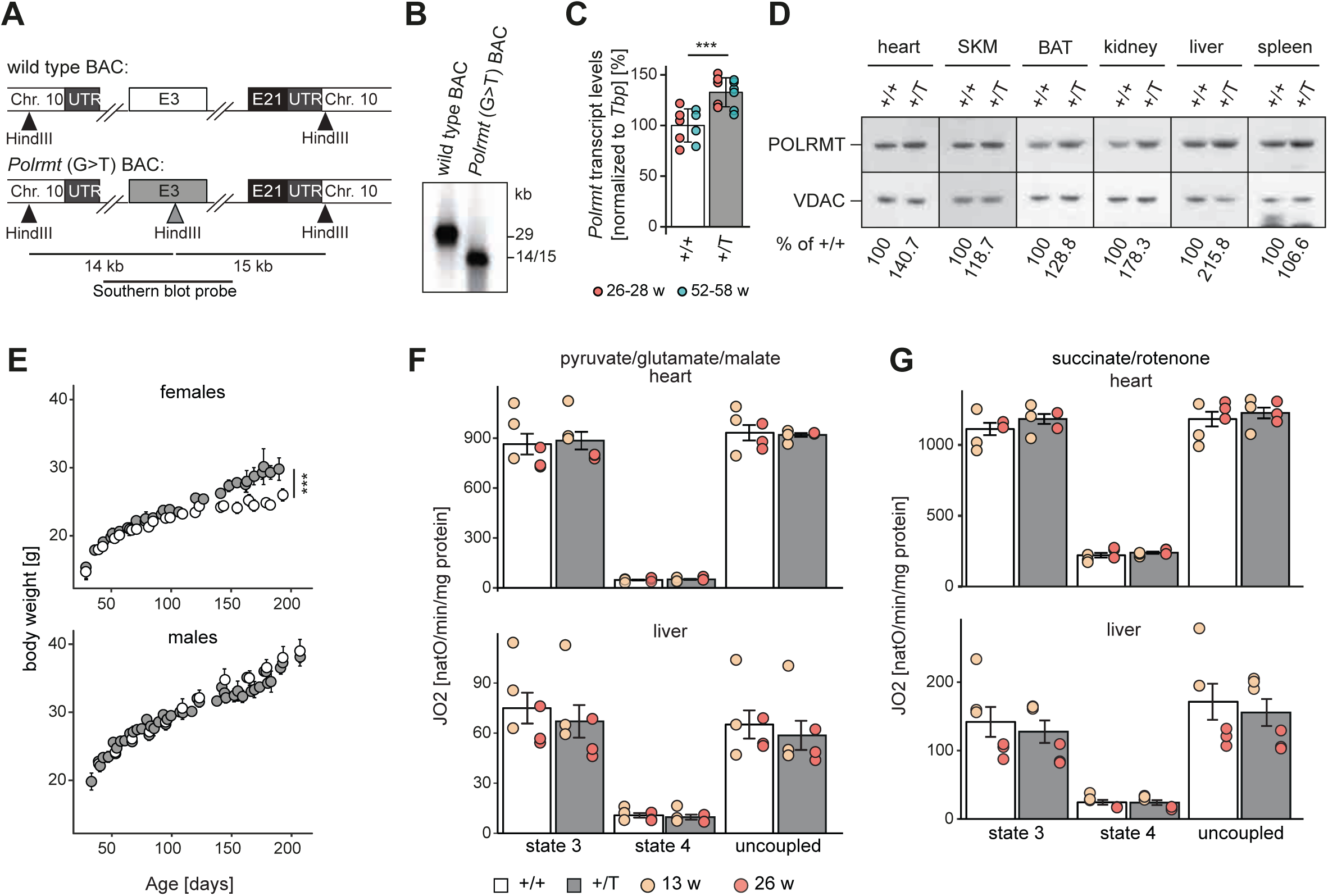
*Polrmt* overexpressing mice are viable. (**A**) Scheme of BAC modification strategy. Chr, chromosome; E, exons; grey triangle indicates introduced HindIII restriction site and black triangles indicate endogenous restriction sites. Size of the DNA fragments generated after HindIII restriction and location of southern blot are indicated at the bottom. (**B**) Representative Southern blot of BAC construct after HindIII restriction digest. (**C**) qRT-PCR analysis of steady-state *Polrmt* transcript levels in wild type (+/+) and *Polrmt* overexpressor (+/T) mouse hearts at different ages. Normalization: tata-binding protein (*Tbp*); n:4-6 per genotype per age). (**D**) Western blot of POLRMT levels in mitochondrial extracts from different tissues of +/+ and +/T 52-week old mice. Loading: VDAC; SKM, skeletal muscle; BAT, brown adipose tissue. n:1 (**E**) Body weight curves of female (upper) and male mice (lower); n: 7-16 per sex and genotype. g, grams; grey:+/T, white:+/+; ***p<0.001 ANOVA repeated measurements (**F-G**) Oxygen consumption analysis on isolated mitochondria from heart and liver. Mitochondria were incubated with pyruvate, glutamate, and malate to deliver electrons to complex I (**F**) or with succinate and rotenone to deliver electrons to complex II (CII) (**G**). Mitochondrial respiration was analyzed under phosphorylating (state 3), non-phosphorylating (state 4), and uncoupled conditions. n: 3 per age. Percentage (%) is calculated relative to +/+ levels. Error bars ± sem. *\*\*\**p<0.001; ANOVA.

### Moderately elevated POLRMT levels have a positive effect on exercise capacity

We assessed the effect of elevated POLRMT levels on animal physiology at different ages, namely at 10, 26 and 52 weeks. Using metabolic cages, we monitored the energy homeostasis and activity of POLRMT overexpressing mice and wild-type littermates during the light and dark cycle. We did not detect significant differences in the drinking and feeding behavior or body weight (Fig S4A-C), which showed that the overexpressors were stress-free when kept under healthy metabolic conditions. We determined the respiratory exchange rate by measuring the O2 consumption and CO2 production and found no differences in substrate utilization *in vivo* between wild-type and POLRMT overexpressor mice (Fig 2A). Surprisingly, when we studied the cumulative distance traveled in the metabolic chambers we observed that POLRMT overexpressing mice tended to display increased activity compared to control mice (Fig 2B). This increase was significant in males at 26 weeks of age in the dark cycle, when mice are more active. We proceeded to study if elevated POLRMT levels could be beneficial for exercise performance in mouse. Using monitored free running wheels and several treadmill runs we challenged the POLRMT overexpressing and littermate control mice of both genders to a voluntary and strenuous exercise regime (Fig 2C). Remarkably, voluntary exercise in the POLRMT overexpressors significantly increased the capacity on a treadmill test compared to controls (Fig 2D, E). The increased distance run on the treadmill test at 10-12 weeks of age and the increased activity in 26 weeks-old male mice show that moderate POLRMT overexpression increases exercise capacity in mice.

**Figure 2:**
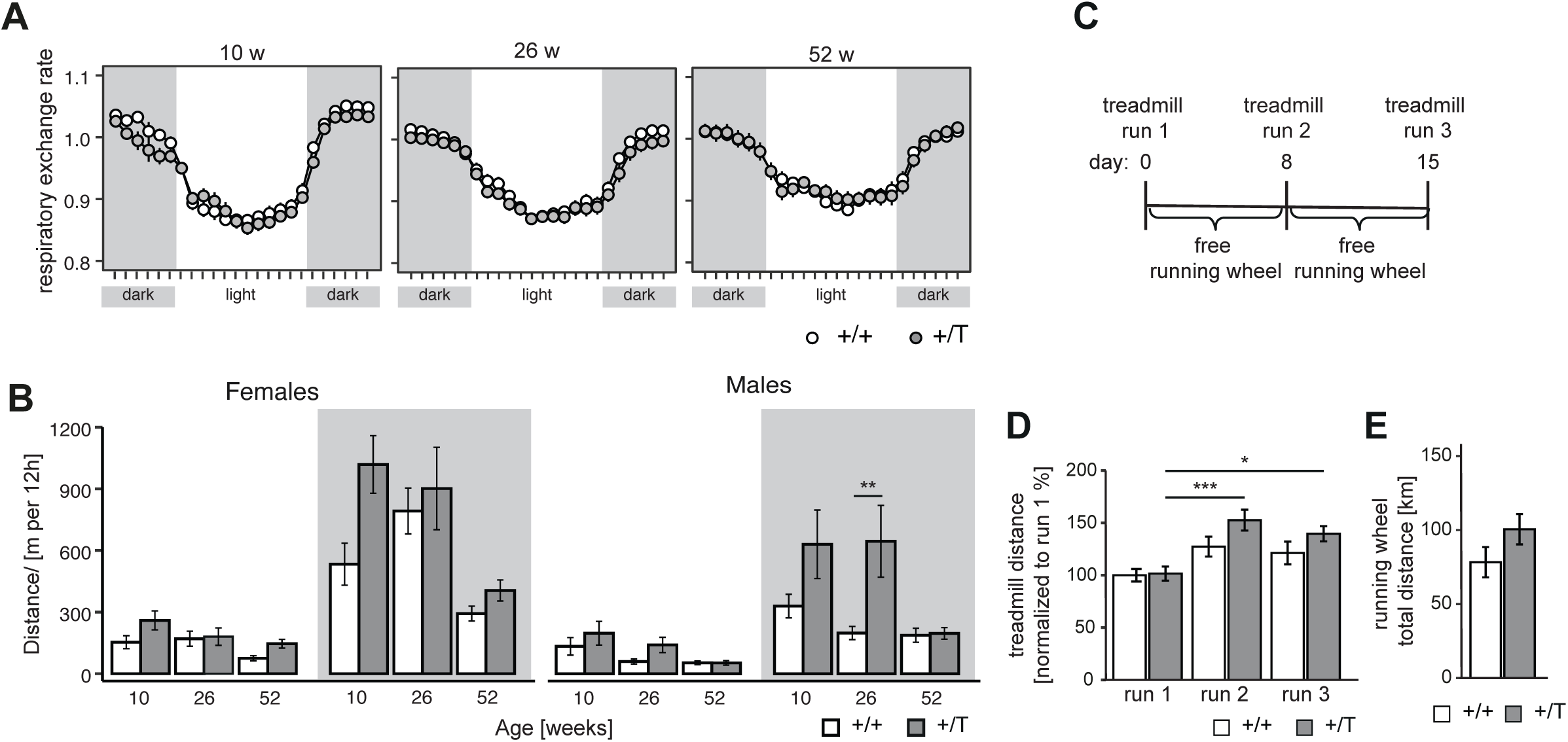
*Polrmt* overexpressing mice have increased exercise capacity. (**A**) Respiratory exchange rate normalized to lean body mass in female and male +/+ and +/T mice at different ages measured in metabolic cages; ticks in x-axis are spaced by 1 h; grey background, dark cycle; white background, light cycle; n:7-9 mice/genotype (**B**) Cumulative distance travelled per day in +/+ and +/T mice at different ages; grey background, dark cycle; white background, light cycle; n:7-9 mice/genotype. (**C**) Scheme exercise challenge on 10-12 week-old females and males +/+ and +/T mice. (**D**) Distance run on treadmill challenge at day 0, 8, and 15. n: 8-12 per genotype. (**E**) Distance run in free running wheels during the exercise challenge. n: 8-12 per genotype. Error bars ± sem. *p<0.05, ***p<0.001, ANOVA.

### POLRMT overexpression moderately increases mtDNA levels

Given the essential role of POLRMT in RNA primer formation, we evaluated whether POLRMT overexpression affected mtDNA replication. We determined steady-state mtDNA copy number by Southern blots and found a mild but significant increase in mtDNA levels (∼17%) in heart of POLRMT overexpressing mice at 26 weeks of age (Fig 3A, B). *De novo* mtDNA synthesis in isolated mitochondria showed a consistent trend to be increased, but this difference was not statistically significant (Fig 3C, D). To evaluate if this increase in mtDNA levels was due to elevated levels of mtDNA maintenance and/or *de novo* replication machinery, we determined protein levels of essential factors for these processes. The protein levels of TFAM (mtDNA maintenance), the mitochondrial DNA helicase TWINKLE and SSBP1 (both for mitochondrial replication) were not changed in POLRMT overexpressing mice (Fig 3E). Our data show that a moderate elevation of POLRMT levels increases mtDNA levels, whereas TFAM levels are unaltered. This finding argues that the TFAM to mtDNA ratio is slightly decreased in POLRMT overexpressing mice.

**Figure 3:**
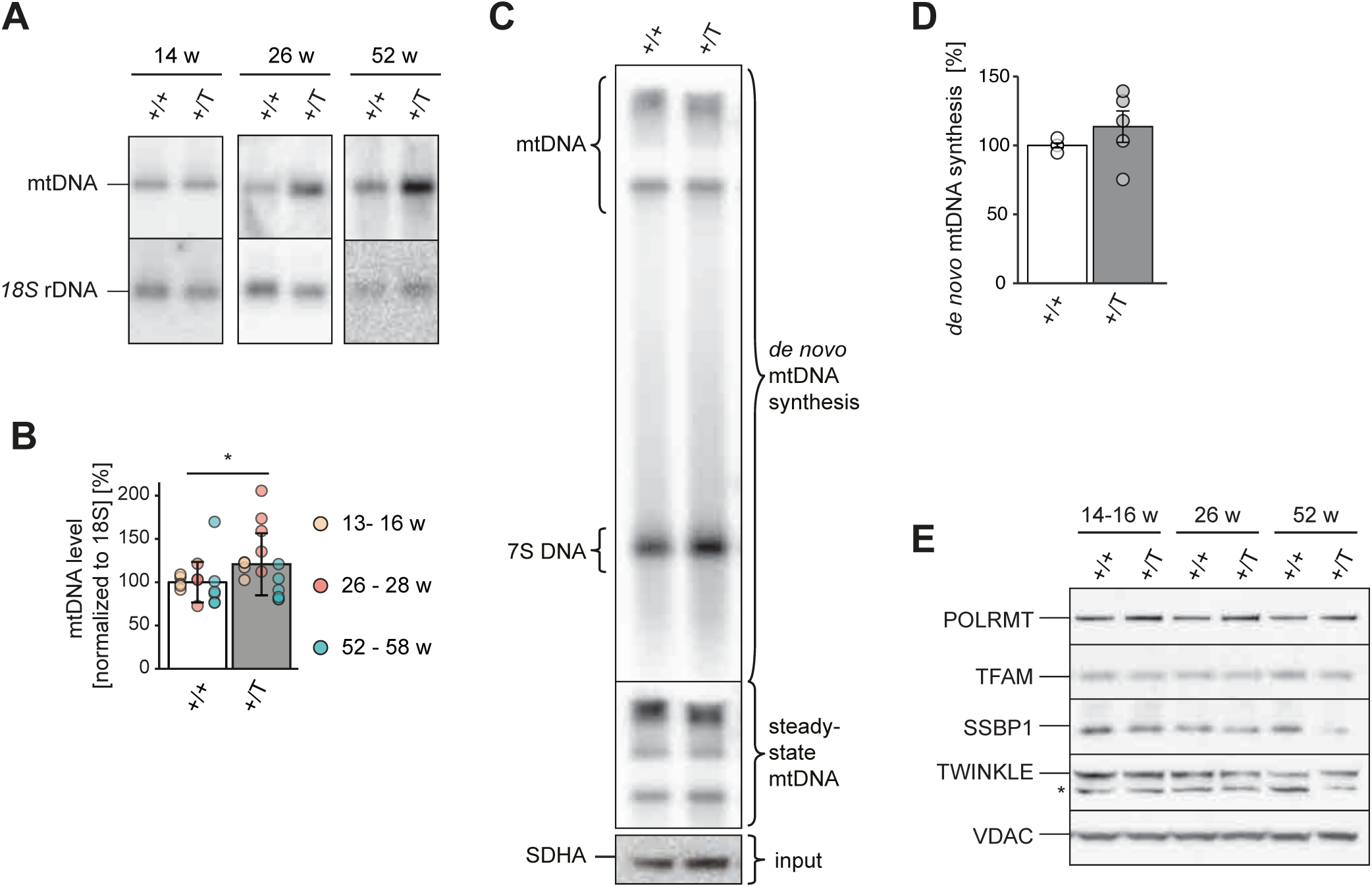
*Polrmt* overexpression increases mtDNA copy number. (**A**) Representative Southern blot analysis of mtDNA levels at different ages. (**B**) Quantification of mtDNA levels from Southern blot at different ages in wild-type (+/+, white) and *Polrmt* overexpressor (+/T, grey) mice. Normalization *18S* rDNA. *p<0.05 ANOVA; n: 4-6. (**C**) Representative *in organello* replication on isolated mitochondria from heart and (**D**) quantification of 3 experiments. Input: Western blot of SHDA and steady-state mtDNA levels. Normalization: protein input; n: 5 per genotype. Error bars ± sem; grey:+/T, white:+/+ (**E**) Western blot of factors required for mtDNA replication expression in +/+ and +/T mice. Loading: VDAC; n:3-4 per genotype; asterisk: unspecific band (Kühl et al., 2016).

### POLRMT levels are limiting mitochondrial transcription *in vivo*

Next, we studied the effect of POLRMT overexpression on mtDNA transcription in mouse hearts. First, we assessed *de novo* transcription in isolated heart mitochondria from POLRMT overexpressing mice and found a significant strong increase of about 50% (Fig 4A, B). There was no accumulation of specific transcription products indicating that the increase in *de novo* transcription is homogeneous and that there is normal processing of the polycistronic mt-RNAs. This finding is consistent with reports that protein-protein interactions between TEFM and RNA processing enzymes may link transcription elongation to RNA processing (Jiang et al., 2019). To assess the steady-state transcript levels of all mt-mRNAs and mt-rRNAs we performed RNA-Sequencing (RNA-Seq) and found similar transcript levels in wild-type and POLRMT overexpressing mice (Fig 4C) despite the strong increase in *de novo* transcription in the POLRMT overexpressing mice. Interestingly, the POLRMT overexpressing mice showed a trend towards slightly elevated levels of the non-polyadenyated *mt-ND6* transcript of the L-strand(Ruzzenente et al., 2012). Next, we investigated whether this discrepancy was accompanied by an imbalance in the protein levels of factors essential for post-transcriptional processes, which could explain why the increased *de novo* transcription does not result in elevated steady-state levels of mt-mRNAs. The steady-state protein levels of factors that are required for mtDNA transcription (TFAM, TEFM, TFB2M), mt-RNA processing and stability (G-rich sequence factor 1, GRSF1, Zinc phosphodiesterase ELAC protein 2, ELAC2, LRPPRC, and SRA stem-loop-interacting RNA-binding protein, SLIRP), and translation (mitochondrial ribosomal proteins, MRPL12, MRPL37, and MRPS35) were not increased despite increased POLRMT levels (Fig 4D). Thus, in mouse hearts increasing POLRMT levels upregulate *de novo* mtDNA transcription (Fig. 4A,B), whereas the steady-state mt-RNA levels are not increased (Fig. 4C).

**Figure 4:**
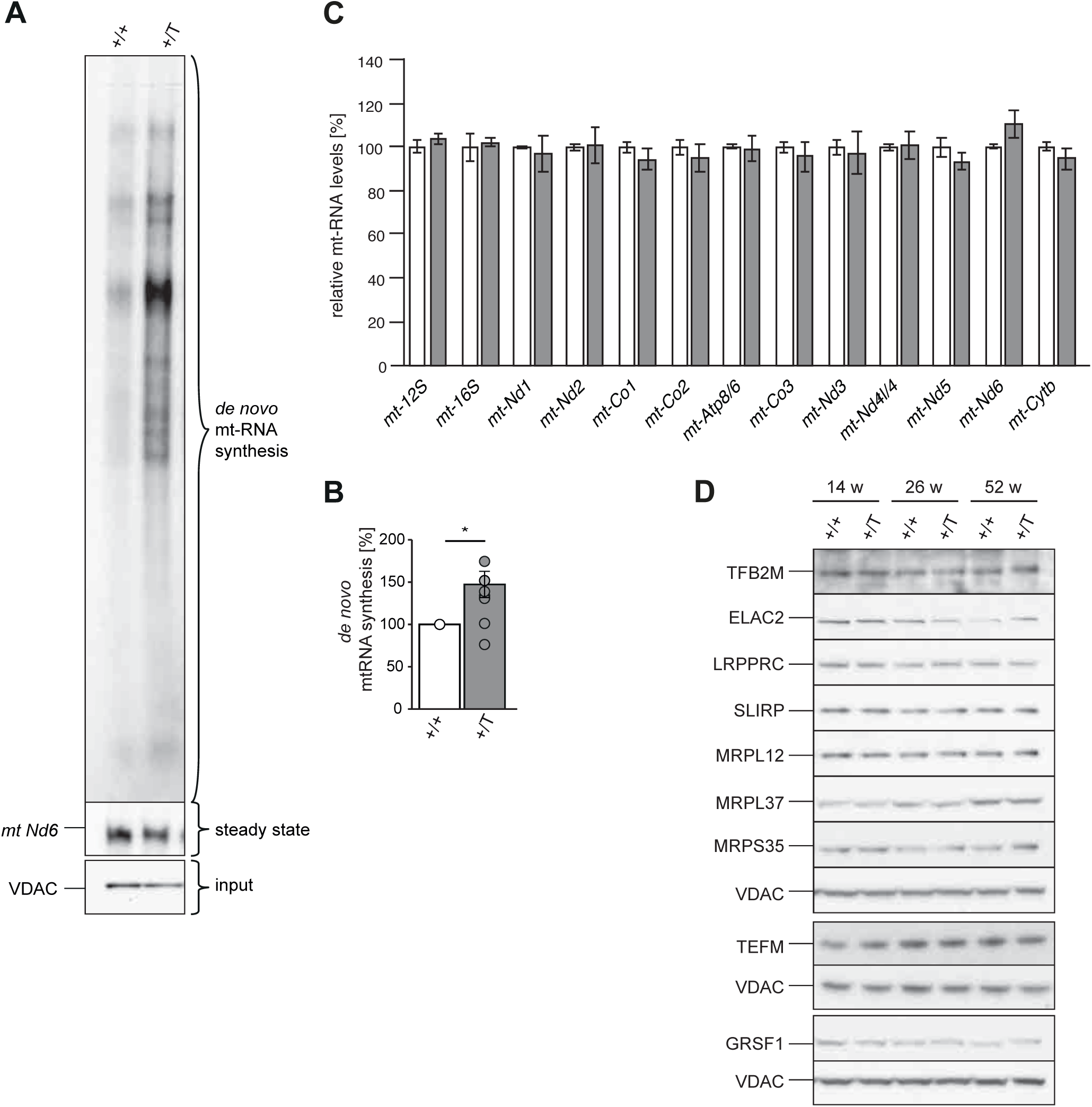
*Polrmt* overexpression increases transcription capacity. (**A**) Representative *de novo* synthesized mitochondrial transcripts analysis from hearts of 14- week old wild-type (+/+) and *Polrmt* overexpressor (+/T) mice. (**B**) Quantification of *de novo* synthesized mitochondrial transcripts normalized to VDAC and +/+ signal; n: 10; *p<0.05, one-sample t-test (μ=100). (**C**) RNA-Seq of mt-rRNAs and mt-mRNAs on total RNA from heart of 14-week old +/+ and +/T mice. n: 3 per genotype. (**D**) Western blot of factors required for mitochondrial transcription or mt-RNA processing and in +/+ and +/T mice. Loading: VDAC; n:3-4. GRSF1 was blotted in the same blot presented in Figure 3E, so VDAC is the same in both figures.

### *Polrmt* overexpression increases *7S* RNA

We next tested whether the discrepancy between the normal steady-state transcript levels and the increased *de novo* transcript synthesis in the POLRMT overexpressing mice could be explained by decreased transcript stability. We followed the relative degradation rate of *de novo* labeled transcripts and found no evidence of increased RNA degradation in the POLRMT overexpressing mitochondria (Fig 5A, B). We quantified the steady-state transcript levels of mt-tRNAs, the precursor transcript *mtNd5/CytB,* and the most promoter-proximal transcript generated from LSP, 7S RNA. While most transcripts remained unchanged in the POLRMT overexpressing mice, we detected a mild decrease in the most promoter-distant LSP transcript *mt-Tq* (Fig 5C, D). On the contrary, we found a striking increase in the most promoter-proximal LSP transcript *7S* RNA across all the time points analyzed in heart (Fig 5C, D) and other mouse tissues (Fig 5E). Increased levels of *7S* RNA have previously been reported to correlate with increased transcription initiation(Cámara et al., 2011; Jiang et al., 2019), and our findings are therefore consistent with highly increased transcription initiation at LSP in the POLRMT overexpressing mice.

**Figure 5:**
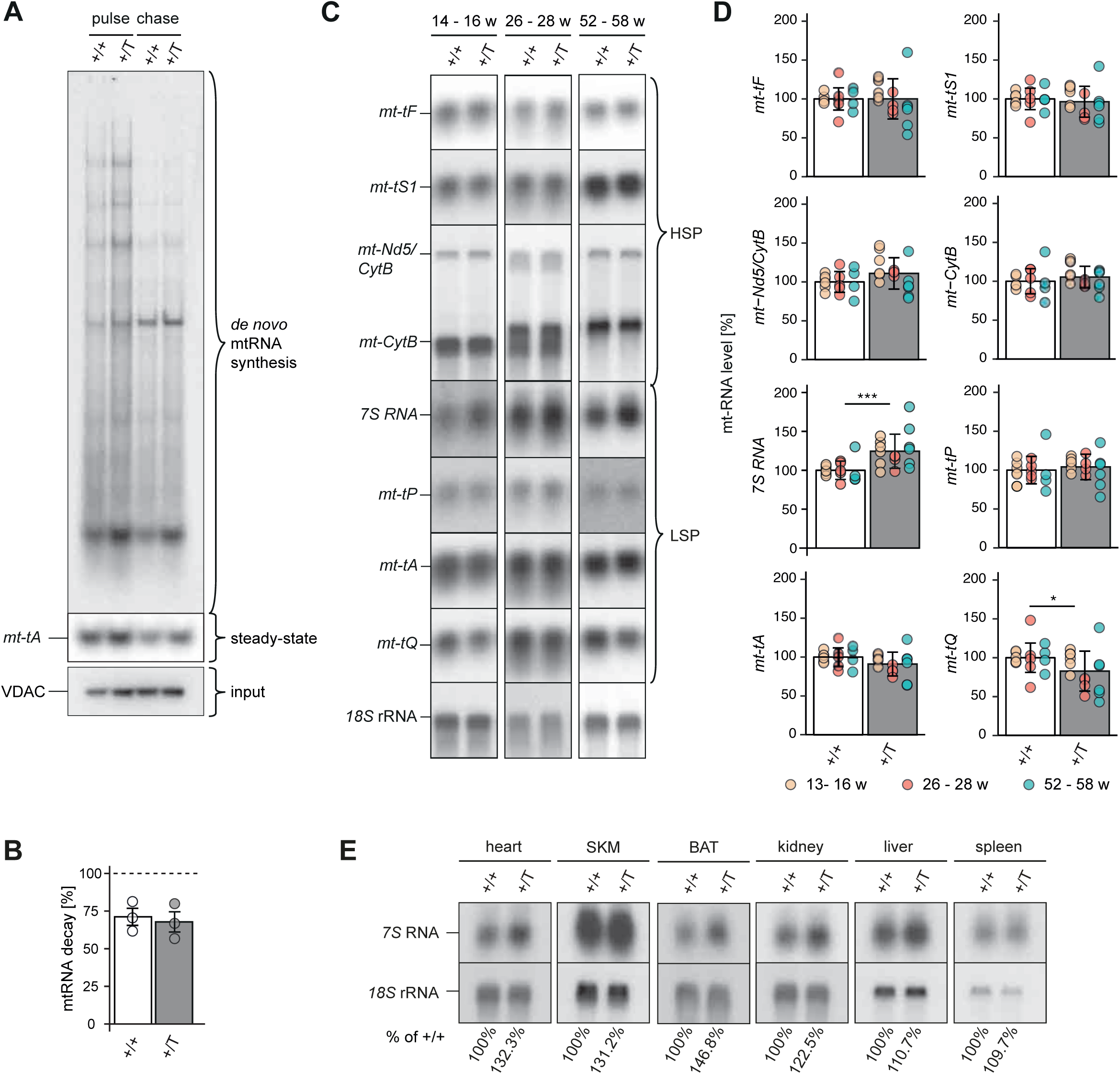
*Polrmt* overexpression increases LSP-promoter proximal 7S RNA. (**A**) *De novo* synthesized mitochondrial transcripts from hearts of 26-week old wild type (+/+) and *Polrmt* overexpressor (+/T) mice (pulse). The mRNA decay of *de novo* synthesized transcripts was followed after 2h (chase). Input: western blot analysis VDAC on radiolabeled mitochondrial extracts. (**B**) Quantification of *de novo* synthesized mitochondrial transcripts normalized to VDAC and pulse signal; n: 3; grey:+/T, white:+/+. (**C**) Representative northern blot analysis of mt-RNA levels in heart of wild-type (+/+) and *Polrmt* overexpressor (+/T) mice at different ages. (**D**) Quantification of mt-RNA levels from northern blot analyses; normalization *18S* rRNA; *p<0.05; ***p<0.001; ANOVA; n: 4-6 per age. (**E**) Northern blot analysis of *7S* RNA levels in different tissues of a +/+ and +/T 52-week old mouse. n:1 per genotype.

### Co-overexpression of LRPPRC and POLRMT does not increase OXPHOS capacity

Since protein levels of the other factors involved in mt-RNA metabolism were unchanged in the POLRMT overexpressing mice (Fig 4D), we hypothesized that the excess of transcripts produced could not be stabilized in the POLRMT overexpressing mice due to limited levels of mt-RNA-stabilizing proteins such as LRPPRC. Therefore, we generated a mouse ubiquitously co-overexpressing LRPPRC^37^ and POLRMT (Fig 6A). LRPPRC overexpression increased steady-state mitochondrial transcripts without affecting OXPHOS capacity as previously shown(Harmel et al., 2013) (Fig 6B, C). The double overexpression of LRPPRC and POLRMT did not show any further increase in the steady-state transcript levels than LRPPRC overexpression alone, and, consistently, OXPHOS protein levels remain normal (Fig 6C). Thus, our data support that the regulatory control of OXPHOS gene expression occurs downstream of transcription initiation and transcript stabilization.

**Figure 6:**
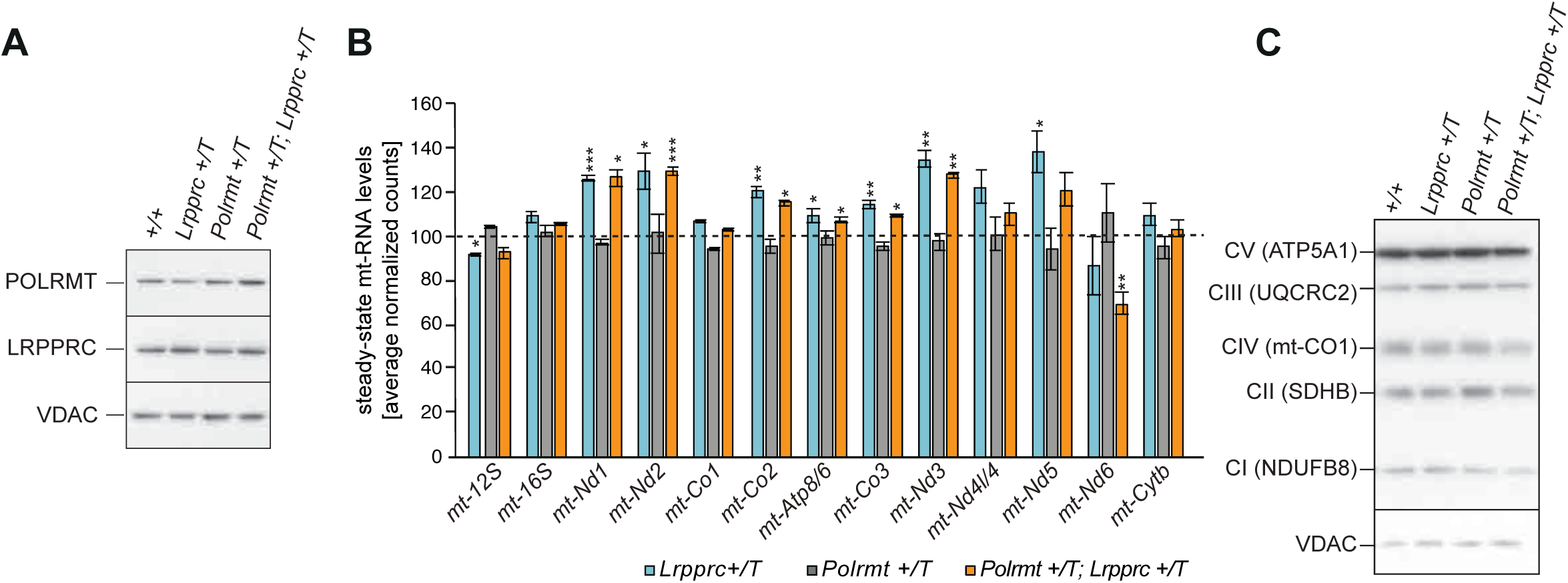
Stabilization of mitochondrial transcripts with *Lrpprc* overexpression does not result in a further increase of mt-RNAs in the *Polrmt* overexpressor mice. **A**) Western blot of steady-state POLRMT and LRPPRC levels in mitochondrial extracts from heart of wild-type (+/+), *Lrpprc* overexpressor (*Lrpprc* +/T), *Polrmt* overexpressor (*Polrmt* +/T), and *Polrmt* and *Lrpprc double* overexpressor (*Polrmt +/T*; *Lrpprc* +/T) mice. Loading: VDAC. (**B**) RNA-Seq of mt-rRNAs and mt-mRNAs on total RNA from heart of 14-week old *Lrpprc* +/T*, Polrmt* +/T, and *Polrmt +/T*; *Lrpprc* +/T. Data is normalized to +/+, represented by the dotted line; n: 3 per genotype. (**C**) Representative western blot of OXPHOS subunit levels in isolated mitochondria from heart at different ages; loading: VDAC. n:3 per genotype.

## DISCUSSION

In this study, we generated and characterized a mouse model overexpressing POLRMT to test whether increasing POLRMT under physiologic conditions modulates OXPHOS function and its effect on health. POLRMT is an essential mitochondrial protein required to initiate mtDNA replication and to transcribe the whole mitochondrial genome. Isolated mitochondria from the POLRMT overexpressing mice had higher levels of *de novo* transcription, indicating that elevated POLRMT levels can increase mitochondrial transcription initiation. Since the protein levels of the other transcription factors were not changed in the POLRMT overexpressing mice, the detection of elevated levels of *de novo* transcripts suggests that a 50% increase of POLRMT is sufficient to upregulate mtDNA transcription and, therefore, we propose that POLRMT is the limiting factor for transcription in mammalian mitochondria. Our data conclusively shows that a moderate 50% increase in POLRMT levels does not affect mitochondrial OXPHOS capacity, is not pathogenic, and does not lead to tumorigenesis until 52 weeks of age, the latest timepoint that we assessed. Only a handful of mitochondrial factors have been proposed to interact with POLRMT (Miranda et al., 2022), but none of them were found to be increased in different tissues of POLRMT overexpressing mice, showing that their expression is not affected by moderately increased POLRMT levels. This finding raises the question why POLRMT is upregulated under several pathogenic conditions.

Transcription initiation from LSP generates the *7S* RNA, a short RNA replication primer, and the full-length polycistronic transcript to express LSP-encoded mitochondrial genes. The robust increase in steady-state levels of mtDNA and *7S* RNA in the POLRMT overexpressing mice argues that transcription initiation at LSP is increased. Although the protein levels of the components of the replisome are unchanged in the POLRMT overexpressing mice, they may be present in excess and may, therefore, not be limiting for mtDNA replication (Jiang et al., 2021). It would be interesting to co-overexpress factors of the replisome in the POLRMT overexpressing background because TWINKLE can be loaded at the 3′ end of 7S DNA to promote full-length genomic replication and has been suggested to play a regulatory role in mtDNA replication (Milenkovic et al., 2013). Moreover, the TFAM to mtDNA ratio is slightly decreased in the POLRMT overexpressor mice because the mtDNA levels increase while TFAM levels remain unchanged. Numerous studies have reported that the amount of TFAM directly regulates mtDNA copy number and that mtDNA levels also reciprocally affect TFAM levels (Bonekamp et al., 2021; Bonekamp and Larsson, 2018). The TFAM to mtDNA ratio has emerged as an important regulator of mitochondrial gene expression, facilitating a switch of nucleoids from an “on” to an “off” state to regulate the number of mtDNA molecules engaged in gene expression (Brüser et al., 2021; Farge et al., 2014). The decreased TFAM to mtDNA ratio in the POLRMT overexpressing mice points to a less compacted mtDNA which is transcribed more when POLRMT levels are increased. POLRMT could thus have a regulatory role in this process.

The discrepancy between increased *de novo* transcription in isolated mitochondria and the unchanged steady-state transcript levels *in vivo* could be explained by the lack of stabilization of the excess transcripts generated *in vivo*. POLRMT produces two near-genomic length polycistronic transcripts that are processed and stabilized co-transcriptionally to generate the mature mt-RNAs. The stability of mt-rRNAs, mt-mRNAs, and mt-tRNAs, is mediated by different sets of proteins; for example, the LRPPRC-SLIRP complex promotes polyadenylation and stabilization of all mt-mRNAs, except *mt-Nd6* (Ruzzenente et al., 2012), whereas mitochondrial ribosomal proteins and assembly factors, which are generally synthesized in excess, stabilize mt-rRNAs (Bogenhagen et al., 2018; Kühl et al., 2017; Park et al., 2007). Increased transcription can therefore lead to different expression patterns of mtDNA encoded genes depending on posttranscriptional events, as exemplified by the *Lrpprc* knockout mice where mt-rRNAs and mt-tRNAs are increased, but mt-mRNAs are depleted (Ruzzenente et al., 2012). None of the mt-rRNAs, mt-tRNAs, or mt-mRNAs are significantly increased on a steady-state level in the POLRMT overexpressor mouse, it is thus unlikely that a moderate POLRMT overexpression leads to a global increase in mitochondrial transcription which cannot be stabilized under normal conditions in vivo. Furthermore, co-overexpressing LRPPRC and POLRMT did not increase mt-mRNAs more than LRPPRC overexpression alone arguing against the hypothesis that transcripts are generated in excess and not stabilized when POLRMT is overexpressed. Although our data does not exclude that protein levels of other factors acting at later steps of mitochondrial transcription stabilization are important, we did not find evidence of decreased transcript stability using pulse-chase *de novo* transcription assays.

An alternative hypothesis to explain the discrepancy between increased *de novo* transcription in isolated mitochondria and the unchanged steady-state transcript is that different transcription scenarios are occurring *in vivo* compared to isolated mitochondria. A strong overexpression of mitochondrial RNA polymerase (Rpo41) in fission yeast increased the mitochondrial transcription capacity and led to increased steady-state levels of mitochondrial transcripts, and was shown to promote the survival of yeast cells from colonies that were exposed to cold (Jiang et al., 2011; Kelly and Lehman, 1986; Masters et al., 1987). Furthermore, previous studies in human cancer cells reported that overexpression of POLRMT from a lentiviral construct (10 fold on mRNA and 2-3 fold increase on protein level (Zhou et al., 2021) resulted in a 2-3 fold increase of mature mt-mRNAs. Importantly, both yeast and cancer cells are actively proliferating cellular models. Despite repeated efforts, we never obtained a founder mouse with more than one additional *Polrm*t gene inserted into the genome. A recent *in vitro* study in cultured human cells has shown that the *7S* RNA has a regulatory role in mitochondrial transcription by directly targeting and blocking the ability of POLRMT to initiate transcription (Zhu et al., 2022). *7S* RNA interacts with POLRMT leading to POLRMT dimerization, which sequesters essential domains for promoter recognition and unwinding. The mitochondrial exoribonuclease complex (mtEXO) is proposed to be directly involved in the degradation of *7S* RNA to overcome the blockage of the pre-initiation complex. Our data is consistent with this mechanism, where the high levels of *7S* RNA may be preventing the increased POLRMT molecules from engaging in full-length mtDNA transcription; thus, most transcripts remain unchanged or slightly decreased, as is the case of *mt-Tq*, under basal physiologic conditions. The increased *de novo* transcription in *in organello* assays could be explained by a release of the negative regulation of *7S* RNA in mitochondrial transcription in response to the ADP-stimulated respiration conditions in which this assay is performed. This raises the question of whether the increased capacity to upregulate mitochondrial transcription could underlie the increased exercise capacity in mice when challenged.

POLRMT is a key factor for mtDNA expression and replication and, thus, essential for OXPHOS biogenesis in mammalian cells. Using several mouse models, we characterized the molecular consequences of ubiquitous POLRMT overexpression in different tissues. We show for the first time that POLRMT is the limiting factor for mtDNA transcription initiation in vivo, but that additional regulatory steps downstream of transcription initiation and transcript stability limit OXPHOS biogenesis. Our data supports a model where POLRMT increases transcription initiation, allowing rapid adaptation to changes in energetic demand, such as increased exercise and, potentially, in some pathogenic states.

## MATERIAL AND METHODS

### Generation and handling of transgenic *Polrmt* overexpressing mice

BAC clones containing a fragment of chromosome 10, including the entire *Polrmt* gene, were purchased from the C57Bl/6N BAC library of DNA Bank, RIKEN BioResource Center. The BAC clone BgN01-092D16 was modified by Red/ET recombination using the counter-selection BAC modification kit (Genebridges). A silent point mutation 420G>T was introduced into exon 3 leading to a unique HindIII restriction site. Positive clones were verified by PCR followed by HindIII restriction digest, Sanger sequencing, and Southern blotting. The modified BAC was purified via a cesium chloride gradient and injected into the pronucleus of fertilized oocytes as described in(Milenkovic et al., 2013). Founders (+/BAC-*Polrmt*) were identified by PCR and restriction enzyme analysis of genomic DNA to detect gain of the HindlII site in the *Polrmt* gene. Tail DNA from offspring was genotyped for the presence of the BAC transgene by analyzing 100 ng of tail DNA with the GoTaq PCR reaction kit (Promega) according to the manufacturer’s instruction by adding forward primer 5′-GAGGCTCGGGTGCGGCAGCTC −3′ and reverse primer 5′-GTGCAGTGTGAGCACCTGCTGTC-3′ for PCR with an initial denaturation for 3 min at 95 °C, followed by 40 cycles for 30 s at 95 °C, 30 s at 60 °C, and 45 s at 72 °C. The reaction was ended with extension for 10 min at 72 °C. Mice were maintained heterozygous on an inbred C57BL/6N background.

### Ethical statement

Animals were housed in individually ventilated cages under specific-pathogen-free conditions with constant temperature (21°C) and humidity (50 - 60%) and a 12 h light/dark cycle. All mice were fed commercial rodent chow and provided with acidified water ad libitum. Mice were sacrificed by cervical dislocation. The health status of the animals is specific pathogen free according to the Federation of the European Laboratory Animal Science Association (FELASA) recommendations. All animal procedures were conducted in accordance with European, national and institutional guidelines and protocols (no.: AZ.: 84-02.05.50.15.004, AZ.: 84- 02.04.2015.A103 and 84-02.04.2016.A420) were approved by the Landesamt für Natur, Umwelt und Verbraucherschutz, Nordrhein-Westfalen, Germany.

### Exercise challenges and phenotyping protocol

Treadmill: Mice were placed on a treadmill (TSE Systems) for a five-minute habituation. After a ten-minute warm-up phase at 0.1 m/sec, speed increased continuously by 0.02 m/min. If mice did not keep up with the treadmill speed, they were exposed to a mild electric shock (0.3 mA). The distance was recorded until mice received three consecutive shocks.

Indirect calorimetry: Indirect calorimetry and home cage locomotor activity were monitored for singly housed mice in purpose built cages (Phenomaster, TSE Systems). Parameters were measured for 48 h after at least four days of acclimatization. For the calculation of the respiratory exchange ratio, the volume of consumed O2 and produced CO2 were normalized to lean body weight.

Running wheel: Voluntary running was monitored for two weeks in individually housed mice using wireless mouse running wheels (Med Associates Inc.).

Body composition: Body fat and lean mass content were measured *in vivo* by nuclear magnetic resonance using the minispec LF50H (Bruker Optics).

### DNA isolation, Southern blot analysis and mtDNA quantification

For genotyping and pyrosequencing, tail or ear-clip biopsies DNA was extracted using chloroform and ethanol precipitation. For Southern blotting or qPCR total DNA was isolated from mouse tissues using the DNeasy Blood & Tissue Kit (QIAGEN) as previously described(Kühl et al., 2016). Briefly, snap frozen tissues were grinded in a cold mortar. About 20 μg of grinded tissue were used to extract DNA using the blood and tissue kit or the Gentra Puregene tissue kit (Qiagen) following manufacturer’s instructions. All samples used for Southern blotting and qPCR were treated with RNase (Qiagen). For Southern blot analysis total DNA (3-10 μg) was digested with SacI endonuclease, fragments were separated by agarose gel electrophoresis, transferred to nitrocellulose membranes (Hybond-N^+^ membranes, GE Healthcare), and hybridized with α-^32^P-dCTP-labeled probes to detect total mtDNA (pAM1) or nuclear DNA (*18S* rDNA) as loading control.

### Pyrosequencing

Quantification of Polrmt c420G>T was performed on tail DNA using a PyroMark-Q24 pyrosequencer (Qiagen). Allele Quantification assay was developed using PyroMark assay design software v. 2.0 (Qiagen). A single PCR reaction was employed to amplify a 192 bp fragment spanning the c.420G>T mutation site, using a biotinylated primer ([Btn]TCTTGCTTGGCTGCAGGTAG) and a non-biotinylated primer (AGAGGCGCCAAAAGGAAGTT). PCR products were combined with distilled water, PyroMark binding buffer (Qiagen), and 1 μL Streptavidin sepharose TM high performance beads (GE Healthcare). Next, PCR products were purified and denatured using a PyroMark Q24 vacuum workstation (Qiagen). Sequencing was performed with PyroMark Gold Q24 reagents according to manufacturer’s instructions using the sequencing primer (CAAGATCTGGAACAAGAA). Relative allele frequencies were calculated using the PyroMark-Q24 Advance v.3.0.0 sfortare (Qiagen).

### RT-PCR, qRT-PCR, northern blot analysis

RNA was isolated either by using the miRNeasy mini kit (QIAGEN). Reverse transcription PCR (RT-PCR) was carried out after cDNA synthesis using the High Capacity cDNA reverse transcription kit (Applied Biosystems). Real-time quantitative reverse transcription PCR (qRT-PCR) was performed using the Taqman 2x Universal PCR mastermix, No Amperase UNG (Applied Biosystems). The quantity of transcripts was normalized to the tata-binding protein (*Tbp)* as a reference gene transcript. For northern blotting, 1-2 μg of total RNA were denatured in RNA Sample Loading buffer (Sigma), separated on 1.2 or 1.8% formaldehyde-MOPS agarose gels prior transfer onto Hybond-N^+^ membranes (GE Healthcare) overnight. After UV crosslinking, the blots were hybridized with various probes at 42°C or 65°C in RapidHyb buffer (Amersham) and then washed in 2x and 0.2x SSC/0.1% SDS before exposure to film. Mitochondrial probes used for visualization of mt-mRNA and mt-rRNA levels were restriction fragments labeled with α-^32^P-dCTP and a random priming kit (Agilent). Different mitochondrial tRNAs and *7S* RNA were detected using specific oligonucleotides labeled with γ-^32^P-ATP. Radioactive signals were detected by autoradiography.

### *In organello* transcription and replication assays

*In organello* transcription and replication assays were performed on mitochondria isolated from mouse hearts by differential centrifugation as described before {Kuhl:2016gu}. *In organello* transcription assays were carried out as previously reported ^38^. For each *in organello* replication assay 1 mg of purified mitochondria were washed in 1 ml of incubation buffer (10 mM Tris pH 7.4, 25 mM sucrose, 75 mM sorbitol, 100 mM KCl, 10 mM K2HPO4, 50 μM EDTA, 5 mM MgCl2, 10 mM glutamate, 2.5 mM malate, 1 mg/ml BSA and 1 mM ADP) and resuspended in 500 μl incubation buffer supplemented with 50 μM of dCTP, dTTP, dGTP and 20 μCi of α-^32^P-dATP (PerkinElmer) and incubated for 2h at 37°C as reported in ^13^. For chase experiments, radiolabeled mitochondria were washed and incubated for more hours 2 hours in incubation buffer without radioactivity. After incubation, mitochondria were washed three times in 10 mM Tris pH 6.8, 0.15 mM MgCl2 and 10% glycerol. An aliquot of radiolabelled mitochondria was collected for immunoblotting with the VDAC (Millipore) or SDHA (Invitrogen) as a loading control. MtDNA was isolated by phenol/chloroform extractions or by Puregene Core Kit A (QIAGEN) and radiolabeled replicated DNA was analyzed by Southern/ D-loop Southern blotting (described earlier) and visualized by autoradiography. Quantifications of transcript levels were performed using the program Multi Gauge with images generated from a PhospoImager instrument.

### RNA sequencing

RNA was isolated from crude heart mitochondria using the miRNeasy mini kit (QIAGEN) and the concentration, purity, and integrity was confirmed using a BioAnalyser. RNA sequencing libraries were constructed using the Illumina TruSeq Sample Prep Kit. Paired end deep sequencing of the mitochondrial RNAs was performed on an Illumina MiSeq according to the manufacturer’s instructions. RNA-Seq was performed on mitochondrial RNA from three biological replicates of each genotype: wild-type, *Polrmt* overexpressor, *Lrpprc* overexpressor, and *Polrmt and Lrpprc* double overexpressor mice aged 14 weeks. Sequenced reads were aligned to the mouse genome (GRCm38) with HISAT 0.1.6 (--fr --rna-strandness RF, total RNA library) (Kim et al., 2015). Reads that aligned to the mitochondria were extracted and subsequently realigned with spliced alignment disabled, to reflect the un-spliced nature of the mitochondrial transcriptome and prevent the introduction of spurious splice junctions. Gene-specific counts were summarised with featureCounts 1.3.5-p4 (Liao et al., 2014)(-p −s 2 −Q 10, total RNA library) using the Ensembl 75 gene annotation for nuclear-encoded genes, and a modified annotation for mitochondrial genes. The modified annotation contains merged *mt-Atp8*/*mt-Atp6* and *mt-Nd4l*/*mt-Nd4* annotations, to reflect their bicistronic nature. Initially, genes with a zero count in any sample were filtered from the count table (in addition to mt-tRNAs), followed by loess and upper quartile normalization for GC-content and sequencing depth, performed with EDASeq 2.3.2 (Risso et al., 2011). Differential expression analysis was performed in R 3.2.0 with edgeR 3.11.2 using a GLM approach with tagwise dispersion estimates and the offsets generated by EDASeq (Robinson et al., 2010).

### Western blots and antisera

For crude mitochondria isolation mouse tissues, were homogenized using a Potter S homogenizer (Sartorius) in mitochondrial isolation buffer (320 mM sucrose, 1 mM EDTA, and 10 mM Tris-HCl pH 7.4) containing complete protease inhibitor cocktail (Roche) followed by two rounds of differential centrifugation. Isolation of crude mitochondria from skeletal muscle was performed as previously described (Frezza et al., 2007). The protein concentration of protein samples was determined using Bradford reagent (Bio-Rad) and BSA as a standard. Proteins were separated by SDS-PAGE (using 4-12% tris-glycine gels, Invitrogen) and then transferred onto polyvinylidene difluoride membranes (GE Healthcare) using wet tank blotting (25 mM Tris, 192 mM glycine, and 20% ethanol) at 4°C at 400 mAmp for 2h or at 80 mA overnight. Immunoblotting was performed according to standard techniques using enhanced chemiluminescence (Immun-Star HRP Luminol/Enhancer from Bio Rad). The following antibodies were used: Total OXPHOS Rodent WB Antibody Cocktail containing NDUFB8 (Complex I), SDHB (Complex II), mt-COI (CIV), UQRC2 (CIII), and ATP5A1 (CV) (abcam ab110413), SDHA (Invitrogen), TFAM (abcam), VDAC (porin) from Mitoscience, and GRSF1, SSBP1, MRPL12, MRPS35, MRPL37 and ELAC2 from Sigma. Further, polyclonal antisera were used to detect TFB2M, SLIRP, LRPPRC, TWINKLE, TEFM and POLRMT proteins (Jiang et al., 2019; Kühl et al., 2014; Lagouge et al., 2015; Milenkovic et al., 2013; Ruzzenente et al., 2012)

### Bioenergetic determinations

Mitochondrial oxygen consumption flux was measured at 37°C using 65–125 μg of crude mitochondria diluted in 2.1 ml of mitochondrial respiration buffer (120 mM sucrose, 50 mM KCl, 20 mM Tris-HCl, 4 mM KH2PO4, 2 mM MgCl2, 1 mM EGTA, pH 7.2) in an Oxygraph-2k (OROBOROS INSTRUMENTS, Innsbruck, Austria). The oxygen consumption rate was measured using either 10 mM pyruvate, 10 mM glutamate, and 5 mM malate or 10 mM succinate and 10 nM rotenone. Oxygen consumption was assessed in the phosphorylating state with 1 mM ADP (state 3) or non-phosphorylating state by adding 2.5 μg/ml oligomycin (state 4). Respiration was uncoupled by successive addition of carbonyl cyanide m-chlorophenyl hydrazone (CCCP) up to 3 µM to reach maximal respiration.

To measure mitochondrial respiratory chain complex activities 15 to 50 µg of mitochondria were diluted in phosphate buffer (KH2PO4 50 mM, pH 7.4), followed by spectrophotometric analysis of isolated respiratory chain complex activities at 37°C, using a HITACHI UV-3600 spectrophotometer. To follow citrate synthase activity, increase in absorbance at 412 nm was recorded after addition of acetyl-CoA (0.1 mM), oxaloacetate (0.5 mM) and DTNB (0.1 mM). Succinate dehydrogenase (SDH) activity was measured at 600 nm after the addition of 10 mM succinate, 35 µM dichlorphenolindophenol (DCPIP) and 1 mM KCN. NADH dehydrogenase activity was determined at 340 nm after the addition of 0.25 mM NADH, 0.25 mM decylubiquinone and 1 mM KCN and controlling for rotenone sensitivity. Cytochrome *c* oxidase activity was measured by standard TMPD ascorbate/KCN sensitive assays. To assess the ATPase activity, 65 µg/ml frozen isolated mitochondria were incubated at 37°C in triethanolamine 75 mM, MgCl2 2 mM, pH 8.9. Mitochondria were pre-incubated 2 min with alamethicin 10 µg/ml prior to addition of 2 mM of ATP. Samples were removed every 2 min and precipitated in 7% HClO4, 25 mM EDTA (50 µl). Phosphate was quantified by incubating an aliquot in 1 ml molybdate 5.34 mM, ferrous sulfate (28.8 mM) and H2SO4 0.75 N for 2 min. The absorbance was assessed at 600 nm. In parallel, oligomycin (2.5 µg per ml protein) was added to the mitochondrial suspension to determine the oligomycin insensitive ATPase activity. Each activity was normalized to mg protein by using the Lowry-based BioRad protein DC kit. All chemicals were obtained from Sigma Aldrich.

### Statistical Analysis

Each mouse was considered an independent biological replicate (n) and repeated measurements of the same biological replicate were considered technical replicates. Unless indicated otherwise, ≥3 biological replicates of transgenic mouse strain and their respective age-matched control mice were used for all the experiments presented. Statistical analyses for RNA-Seq were performed as described above. For qRT-PCR and metabolomics analysis variance was assessed using an F-test and statistical significance was assessed by a two-sample, two-tailed unpaired Student’s t-test in Excel. *In organello* experiments were analyzed using a one-sample, two-tailed Student’s t-test with μ:100 as each experiment was normalized as percentage of the wild-type control that was run simultaneously. Multiple comparison was corrected using Benjamini-Hochberg or Bonferroni correction as indicated in the figure legends. Statistical analyses were performed in Excel or R Studio version 1.1.383. Data visualization in R was done using ggplot2 version 2.2.1 respectively. The definition of center and precision measures, and p values are provided in the figure legends. p<0.05 was considered significant.

## ACKNOWLEDGEMENTS

We are grateful for technical assistance to Magdalena Springer, Nadine Hochhard, and Lysann Schmitz. We thank James B. Stewart and Laila Singh for assistance with pyrosequencing and Paola Loguercio Polosa for technical advice and fruitful discussions. Transgenic animals were generated with the help of Ingo Voigt, Transgenesis core facility of the Max Planck Institute for Biology of Ageing. RNA library construction and sequencing were performed at the Cologne Center for Genomics.

## Author contributions

Experiments were done by A.Me., A.Mo., I.Kü., L.P., L.S., M.M., M.P.; M.M. performed most of the experiments. A.F. and I.Ku. did the RNA-Seq data analysis. M.M, I.Kü. and N.-G.L. designed and conceived the experiments, integrated all the results, discussed the analysis and wrote the manuscript with contributions from all the co-authors. I.Kü. and N.-G.L. supervised and had the original idea of the project.

## REFERENCES

Agaronyan, K., Morozov, Y.I., Anikin, M., Temiakov, D., 2015. Replication-transcription switch in human mitochondria. Science 347, 548–551. 10.1126/science.aaa0986

Bogenhagen, D.F., Ostermeyer-Fay, A.G., Haley, J.D., Garcia-Diaz, M., 2018. Kinetics and Mechanism of Mammalian Mitochondrial Ribosome Assembly. Cell Rep. 22, 1935– 1944. 10.1016/j.celrep.2018.01.066

Bonekamp, N.A., Jiang, M., Motori, E., Garcia Villegas, R., Koolmeister, C., Atanassov, I., Mesaros, A., Park, C.B., Larsson, N.-G., 2021. High levels of TFAM repress mammalian mitochondrial DNA transcription in vivo. Life Sci. Alliance 4, e202101034. 10.26508/lsa.202101034

Bonekamp, N.A., Larsson, N.-G., 2018. SnapShot: Mitochondrial Nucleoid. Cell 172, 388–388.e1. 10.1016/j.cell.2017.12.039

Bonekamp, N.A., Peter, B., Hillen, H.S., Felser, A., Bergbrede, T., Choidas, A., Horn, M., Unger, A., Di Lucrezia, R., Atanassov, I., Li, X., Koch, U., Menninger, S., Boros, J., Habenberger, P., Giavalisco, P., Cramer, P., Denzel, M.S., Nussbaumer, P., Klebl, B., Falkenberg, M., Gustafsson, C.M., Larsson, N.-G., 2020. Small-molecule inhibitors of human mitochondrial DNA transcription. Nature 588, 712–716. 10.1038/s41586-020-03048-z

Bralha, F.N., Liyanage, S.U., Hurren, R., Wang, X., Son, M.H., Fung, T.A., Chingcuanco, F.B., Tung, A.Y.W., Andreazza, A.C., Psarianos, P., Schimmer, A.D., Salmena, L., Laposa, R.R., 2015. Targeting mitochondrial RNA polymerase in acute myeloid leukemia. Oncotarget 6, 37216–37228. 10.18632/oncotarget.6129

Brüser, C., Keller-Findeisen, J., Jakobs, S., 2021. The TFAM-to-mtDNA ratio defines inner-cellular nucleoid populations with distinct activity levels. Cell Rep. 37, 110000. 10.1016/j.celrep.2021.110000

Cámara, Y., Asin-Cayuela, J., Park, C.B., Metodiev, M.D., Shi, Y., Ruzzenente, B., Kukat, C., Habermann, B., Wibom, R., Hultenby, K., Franz, T., Erdjument-Bromage, H., Tempst, P., Hallberg, B.M., Gustafsson, C.M., Larsson, N.-G., 2011. MTERF4 Regulates Translation by Targeting the Methyltransferase NSUN4 to the Mammalian Mitochondrial Ribosome. Cell Metab. 13, 527–539. 10.1016/j.cmet.2011.04.002

Chang, D., Hauswirth, W., Clayton, D., 1985. Replication priming and transcription initiate from precisely the same site in mouse mitochondrial DNA. EMBO J. 4, 1559–1567.

Chaudhary, S., Ganguly, S., Palanichamy, J.K., Singh, A., Bakhshi, R., Jain, A., Chopra, A., Bakhshi, S., 2021. PGC1A driven enhanced mitochondrial DNA copy number predicts outcome in pediatric acute myeloid leukemia. Mitochondrion 58, 246–254. 10.1016/j.mito.2021.03.013

Crews, S., Ojala, D., Posakony, J., Nishiguchi, J., Attardi, G., 1979. Nucleotide sequence of a region of human mitochondrial DNA containing the precisely identified origin of replication. Nature 277, 192–198. 10.1038/277192a0

Falkenberg, M., 2018. Mitochondrial DNA replication in mammalian cells: overview of the pathway. Essays Biochem. 62, 287–296. 10.1042/EBC20170100

Farge, G., Mehmedovic, M., Baclayon, M., van den Wildenberg, S.M.J.L., Roos, W.H., Gustafsson, C.M., Wuite, G.J.L., Falkenberg, M., 2014. In Vitro-Reconstituted Nucleoids Can Block Mitochondrial DNA Replication and Transcription. Cell Rep. 8, 66–74. 10.1016/j.celrep.2014.05.046

Frezza, C., Cipolat, S., Scorrano, L., 2007. Organelle isolation: functional mitochondria from mouse liver, muscle and cultured filroblasts. Nat. Protoc. 2, 287–295. 10.1038/nprot.2006.478

Gohil, V.M., Nilsson, R., Belcher-Timme, C.A., Luo, B., Root, D.E., Mootha, V.K., 2010. Mitochondrial and Nuclear Genomic Responses to Loss of LRPPRC Expression. J. Biol. Chem. 285, 13742–13747. 10.1074/jbc.M109.098400

Gustafsson, C.M., Falkenberg, M., Larsson, N.-G., 2016. Maintenance and Expression of Mammalian Mitochondrial DNA. Annu. Rev. Biochem. 85, 133–160. 10.1146/annurev-biochem-060815-014402

Harmel, J., Ruzzenente, B., Terzioglu, M., Spåhr, H., Falkenberg, M., Larsson, N.-G., 2013. The Leucine-rich Pentatricopeptide Repeat-containing Protein (LRPPRC) Does Not Activate Transcription in Mammalian Mitochondria. J. Biol. Chem. 288, 15510–15519. 10.1074/jbc.M113.471649

Hillen, H.S., Morozov, Y.I., Sarfallah, A., Temiakov, D., Cramer, P., 2017. Structural Basis of Mitochondrial Transcription Initiation. Cell 171, 1072–1081.e10. 10.1016/j.cell.2017.10.036

Hillen, H.S., Temiakov, D., Cramer, P., 2018. Structural basis of mitochondrial transcription. Nat. Struct. Mol. Biol. 25, 754–765. 10.1038/s41594-018-0122-9

Holzmann, J., Frank, P., Löffler, E., Bennett, K.L., Gerner, C., Rossmanith, W., 2008. RNase P without RNA: Identification and Functional Reconstitution of the Human Mitochondrial tRNA Processing Enzyme. Cell 135, 462–474. 10.1016/j.cell.2008.09.013

Jiang, H., Sun, W., Wang, Z., Zhang, J., Chen, D., Murchie, A.I.H., 2011. Identification and characterization of the mitochondrial RNA polymerase and transcription factor in the fission yeast Schizosaccharomyces pombe. Nucleic Acids Res. 39, 5119–5130. 10.1093/nar/gkr103

Jiang, M., Xie, X., Zhu, X., Jiang, S., Milenkovic, D., Misic, J., Shi, Y., Tandukar, N., Li, X., Atanassov, I., Jenninger, L., Hoberg, E., Albarran-Gutierrez, S., Szilagyi, Z., Macao, B., Siira, S.J., Carelli, V., Griffith, J.D., Gustafsson, C.M., Nicholls, T.J., Filipovska, A., Larsson, N.-G., Falkenberg, M., 2021. The mitochondrial single-stranded DNA binding protein is essential for initiation of mtDNA replication. Sci. Adv. 7, eabf8631. 10.1126/sciadv.abf8631

Jiang, S., Koolmeister, C., Misic, J., Siira, S., Kühl, I., Silva Ramos, E., Miranda, M., Jiang, M., Posse, V., Lytovchenko, O., Atanassov, I., Schober, F.A., Wibom, R., Hultenby, K., Milenkovic, D., Gustafsson, C.M., Filipovska, A., Larsson, N., 2019. TEFM regulates both transcription elongation and RNA processing in mitochondria. EMBO Rep. 20. 10.15252/embr.201948101

Kelly, J., Lehman, I., 1986. Yeast mitochondrial RNA polymerase. Purification and properties of the catalytic subunit. J Biol Chem 261, 10340–7.

Kim, D., Langmead, B., Salzberg, S.L., 2015. HISAT: a fast spliced aligner with low memory requirements. Nat. Methods 12, 357–360. 10.1038/nmeth.3317

Kühl, I., Kukat, C., Ruzzenente, B., Milenkovic, D., Mourier, A., Miranda, M., Koolmeister, C., Falkenberg, M., Larsson, N.-G., 2014. POLRMT does not transcribe nuclear genes. Nature 514, E7–E11. 10.1038/nature13690

Kühl, I., Miranda, M., Atanassov, I., Kuznetsova, I., Hinze, Y., Mourier, A., Filipovska, A., Larsson, N.-G., 2017. Transcriptomic and proteomic landscape of mitochondrial dysfunction reveals secondary coenzyme Q deficiency in mammals. eLife 6, e30952. 10.7554/eLife.30952

Kühl, I., Miranda, M., Posse, V., Milenkovic, D., Mourier, A., Siira, S.J., Bonekamp, N.A., Neumann, U., Filipovska, A., Polosa, P.L., Gustafsson, C.M., Larsson, N.-G., 2016. POLRMT regulates the switch between replication primer formation and gene expression of mammalian mtDNA. Sci. Adv. 2, e1600963. 10.1126/sciadv.1600963

Lagouge, M., Mourier, A., Lee, H.J., Spåhr, H., Wai, T., Kukat, C., Silva Ramos, E., Motori, E., Busch, J.D., Siira, S., German Mouse Clinic Consortium, Kremmer, E., Filipovska, A., Larsson, N.-G., 2015. SLIRP Regulates the Rate of Mitochondrial Protein Synthesis and Protects LRPPRC from Degradation. PLOS Genet. 11, e1005423. 10.1371/journal.pgen.1005423

Liao, Y., Smyth, G.K., Shi, W., 2014. featureCounts: an efficient general purpose program for assigning sequence reads to genomic features. Bioinformatics 30, 923–930. 10.1093/bioinformatics/btt656

Masters, B.S., Stohl, L.L., Clayton, D.A., 1987. Yeast mitochondrial RNA polymerase is homologous to those encoded by bacteriophages T3 and T7. Cell 51, 89–99. 10.1016/0092-8674(87)90013-4

Milenkovic, D., Matic, S., Kuhl, I., Ruzzenente, B., Freyer, C., Jemt, E., Park, C.B., Falkenberg, M., Larsson, N.-G., 2013. TWINKLE is an essential mitochondrial helicase required for synthesis of nascent D-loop strands and complete mtDNA replication. Hum. Mol. Genet. 22, 1983–1993. 10.1093/hmg/ddt051

Minczuk, M., He, J., Duch, A.M., Ettema, T.J., Chlebowski, A., Dzionek, K., Nijtmans, L.G.J., Huynen, M.A., Holt, I.J., 2011. TEFM (c17orf42) is necessary for transcription of human mtDNA. Nucleic Acids Res. 39, 4284–4299. 10.1093/nar/gkq1224

Miranda, M., Bonekamp, N.A., Kühl, I., 2022. Starting the engine of the powerhouse: mitochondrial transcription and beyond. Biol. Chem. 403, 779–805. 10.1515/hsz-2021-0416

Montoya, J., Christianson, T., Levens, D., Rabinowitz, M., Attardi, G., 1982. Identification of initiation sites for heavy-strand and light-strand transcription in human mitochondrial DNA. Proc. Natl. Acad. Sci. 79, 7195–7199. 10.1073/pnas.79.23.7195

Ojala, D., Montoya, J., Attardi, G., 1981. tRNA punctuation model of RNA processing in human mitochondria. Nature 290, 470–474. 10.1038/290470a0

Oláhová, M., Peter, B., Szilagyi, Z., Diaz-Maldonado, H., Singh, M., Sommerville, E.W., Blakely, E.L., Collier, J.J., Hoberg, E., Stránecký, V., Hartmannová, H., Bleyer, A.J., McBride, K.L., Bowden, S.A., Korandová, Z., Pecinová, A., Ropers, H.-H., Kahrizi, K., Najmabadi, H., Tarnopolsky, M.A., Brady, L.I., Weaver, K.N., Prada, C.E., Õunap, K., Wojcik, M.H., Pajusalu, S., Syeda, S.B., Pais, L., Estrella, E.A., Bruels, C.C., Kunkel, L.M., Kang, P.B., Bonnen, P.E., Mráček, T., Kmoch, S., Gorman, G.S., Falkenberg, M., Gustafsson, C.M., Taylor, R.W., 2021. POLRMT mutations impair mitochondrial transcription causing neurological disease. Nat. Commun. 12, 1135. 10.1038/s41467-021-21279-0

Park, C.B., Asin-Cayuela, J., Cámara, Y., Shi, Y., Pellegrini, M., Gaspari, M., Wibom, R., Hultenby, K., Erdjument-Bromage, H., Tempst, P., Falkenberg, M., Gustafsson, C.M., Larsson, N.-G., 2007. MTERF3 Is a Negative Regulator of Mammalian mtDNA Transcription. Cell 130, 273–285. 10.1016/j.cell.2007.05.046

Perks, K.L., Rossetti, G., Kuznetsova, I., Hughes, L.A., Ermer, J.A., Ferreira, N., Busch, J.D., Rudler, D.L., Spahr, H., Schöndorf, T., Shearwood, A.-M.J., Viola, H.M., Siira, S.J., Hool, L.C., Milenkovic, D., Larsson, N.-G., Rackham, O., Filipovska, A., 2018. PTCD1 Is Required for 16S rRNA Maturation Complex Stability and Mitochondrial Ribosome Assembly. Cell Rep. 23, 127–142. 10.1016/j.celrep.2018.03.033

Rackham, O., Busch, J.D., Matic, S., Siira, S.J., Kuznetsova, I., Atanassov, I., Ermer, J.A., Shearwood, A.-M.J., Richman, T.R., Stewart, J.B., Mourier, A., Milenkovic, D., Larsson, N.-G., Filipovska, A., 2016. Hierarchical RNA Processing Is Required for Mitochondrial Ribosome Assembly. Cell Rep. 16, 1874–1890. 10.1016/j.celrep.2016.07.031

Rath, S., Sharma, R., Gupta, R., Ast, T., Chan, C., Durham, T.J., Goodman, R.P., Grabarek, Z., Haas, M.E., Hung, W.H.W., Joshi, P.R., Jourdain, A.A., Kim, S.H., Kotrys, A.V., Lam, S.S., McCoy, J.G., Meisel, J.D., Miranda, M., Panda, A., Patgiri, A., Rogers, R., Sadre, S., Shah, H., Skinner, O.S., To, T.-L., Walker, M.A., Wang, H., Ward, P.S., Wengrod, J., Yuan, C.-C., Calvo, S.E., Mootha, V.K., 2021. MitoCarta3.0: an updated mitochondrial proteome now with sub-organelle localization and pathway annotations. Nucleic Acids Res. 49, D1541–D1547. 10.1093/nar/gkaa1011

Ringel, R., Sologub, M., Morozov, Y.I., Litonin, D., Cramer, P., Temiakov, D., 2011. Structure of human mitochondrial RNA polymerase. Nature 478, 269–273. 10.1038/nature10435

Risso, D., Schwartz, K., Sherlock, G., Dudoit, S., 2011. GC-Content Normalization for RNA-Seq Data. BMC Bioinformatics 12, 480. 10.1186/1471-2105-12-480

Robinson, M.D., McCarthy, D.J., Smyth, G.K., 2010. edgeR: a Bioconductor package for differential expression analysis of digital gene expression data. Bioinformatics 26, 139–140. 10.1093/bioinformatics/btp616

Ruzzenente, B., Metodiev, M.D., Wredenberg, A., Bratic, A., Park, C.B., Cámara, Y., Milenkovic, D., Zickermann, V., Wibom, R., Hultenby, K., Erdjument-Bromage, H., Tempst, P., Brandt, U., Stewart, J.B., Gustafsson, C.M., Larsson, N.-G., 2012. LRPPRC is necessary for polyadenylation and coordination of translation of mitochondrial mRNAs: LRPPRC regulates mitochondrial translation. EMBO J. 31, 443–456. 10.1038/emboj.2011.392

Sarfallah, A., Zamudio-Ochoa, A., Anikin, M., Temiakov, D., 2021. Mechanism of transcription initiation and primer generation at the mitochondrial replication origin OriL. EMBO J. 40. 10.15252/embj.2021107988

Sasarman, F., Brunel-Guitton, C., Antonicka, H., Wai, T., Shoubridge, E.A., LSFC Consortium, 2010. LRPPRC and SLIRP Interact in a Ribonucleoprotein Complex That Regulates Posttranscriptional Gene Expression in Mitochondria. Mol. Biol. Cell 21, 1315–1323. 10.1091/mbc.e10-01-0047

Siira, S.J., Rossetti, G., Richman, T.R., Perks, K., Ermer, J.A., Kuznetsova, I., Hughes, L., Shearwood, A.J., Viola, H.M., Hool, L.C., Rackham, O., Filipovska, A., 2018. Concerted regulation of mitochondrial and nuclear non-coding RNA s by a dual-targeted RN ase Z. EMBO Rep. 19, e46198. 10.15252/embr.201846198

Silva Ramos, E., Motori, E., Brüser, C., Kühl, I., Yeroslaviz, A., Ruzzenente, B., Kauppila, J.H.K., Busch, J.D., Hultenby, K., Habermann, B.H., Jakobs, S., Larsson, N.-G., Mourier, A., 2019. Mitochondrial fusion is required for regulation of mitochondrial DNA replication. PLOS Genet. 15, e1008085. 10.1371/journal.pgen.1008085

Sotgia, F., Whitaker-Menezes, D., Martinez-Outschoorn, U.E., Salem, A.F., Tsirigos, A., Lamb, R., Sneddon, S., Hulit, J., Howell, A., Lisanti, M.P., 2012. Mitochondria “fuel” breast cancer metabolism: Fifteen markers of mitochondrial biogenesis label epithelial cancer cells, but are excluded from adjacent stromal cells. Cell Cycle 11, 4390–4401. 10.4161/cc.22777

Tan, B.G., Mutti, C.D., Shi, Y., Xie, X., Zhu, X., Silva-Pinheiro, P., Menger, K.E., Díaz-Maldonado, H., Wei, W., Nicholls, T.J., Chinnery, P.F., Minczuk, M., Falkenberg, M., Gustafsson, C.M., 2022. The human mitochondrial genome contains a second light strand promoter. Mol. Cell 82, 3646–3660.e9. 10.1016/j.molcel.2022.08.011

Terzioglu, M., Ruzzenente, B., Harmel, J., Mourier, A., Jemt, E., López, M.D., Kukat, C., Stewart, J.B., Wibom, R., Meharg, C., Habermann, B., Falkenberg, M., Gustafsson, C.M., Park, C.B., Larsson, N.-G., 2013. MTERF1 Binds mtDNA to Prevent Transcriptional Interference at the Light-Strand Promoter but Is Dispensable for rRNA Gene Transcription Regulation. Cell Metab. 17, 618–626. 10.1016/j.cmet.2013.03.006

Wanrooij, P.H., Uhler, J.P., Shi, Y., Westerlund, F., Falkenberg, M., Gustafsson, C.M., 2012. A hybrid G-quadruplex structure formed between RNA and DNA explains the extraordinary stability of the mitochondrial R-loop. Nucleic Acids Res. 40, 10334– 10344. 10.1093/nar/gks802

Zhou, T., Sang, Y.-H., Cai, S., Xu, C., Shi, M., 2021. The requirement of mitochondrial RNA polymerase for non-small cell lung cancer cell growth. Cell Death Dis. 12, 751. 10.1038/s41419-021-04039-2

Zhu, X., Xie, X., Das, H., Tan, B.G., Shi, Y., Al-Behadili, A., Peter, B., Motori, E., Valenzuela, S., Posse, V., Gustafsson, C.M., Hällberg, B.M., Falkenberg, M., 2022. Non-coding 7S RNA inhibits transcription via mitochondrial RNA polymerase dimerization. Cell 185, 2309–2323.e24. 10.1016/j.cell.2022.05.006

