## Supplementary material for "Increased mitochondrial transcription initiation does not promote oxidative phosphorylation": FiguresSupplements_Miranda et al_BioRxiv.pdf

### SUPPLEMENTARY FIGURE LEGENDS:

**Supplementary Figure 1: Generation of endogenous *Polrmt* overexpressing mice**
(**A**) Detailed scheme of BAC modification strategy. Chr, chromosome; E, exons; asterisk point mutation introduced in c420. Sequencing chromatogram is shown. (**B**) PCR and restriction digest analysis of the *Polrmt* (G>T) BAC transgenic allele in genomic (gDNA) and reverse transcribed RNA (cDNA) from wild-type (+/+) and BAC transgenic *Polrmt* overexpressor (+/T) mice. (**C**) Pyrosequencing analysis of the ratio of wild-type (G) and transgenic (T) alleles in DNA isolated from tail biopsies in +/+ and +/T mice; n: 9 per genotype and sex.

**Supplementary Figure 2: *Polrmt* overexpressor mice are healthy**
(**A**) Histogram of genotype distribution of the wild-type (+/+, white) and *Polrmt* overexpressor (+/T, grey) offspring; n=22-28 per genotype per sex, n.s: p>0.05; chi-square test. (**B**) Fat mass percentage and (**C**) Lean mass percentage of 10-week-old +/+ and +/T mice before starting the exercise challenge; n: 8-12 per sex per genotype. (**D**) Heart to body weight ratio at different ages; n: 3-5 per age and genotype.

**Supplementary Figure 3: Bioenergetic analyses in heart and liver mitochondria**
(**A**) Relative enzyme activity of respiratory chain enzymes measured in mitochondria isolated from heart and liver at different ages. The enzymes measured are: CI: NADH ubiquinone oxidoreductase, CII: succinate dehydrogenase, Complex II-III: Succinate dehydrogenase - cytochrome c reductase, Complex IV (CIV): Cytochrome c oxidase and citrate synthase (CS). n: 3 per age. (**B**) Western blot of OXPHOS subunit levels in isolated mitochondria from heart at different ages; loading: VDAC. n:3. Error bars ± sem.

**Supplementary Figure 4: Phenomaster parameters**
(**A**) Body weight of female and male wild type (+/+) and *Polrmt* overexpressor (+/T) mice at different ages at the beginning of indirect calorimetry experiment. (**B**) Water consumption. (**C**) Food consumption. n:8-9 mice/genotype

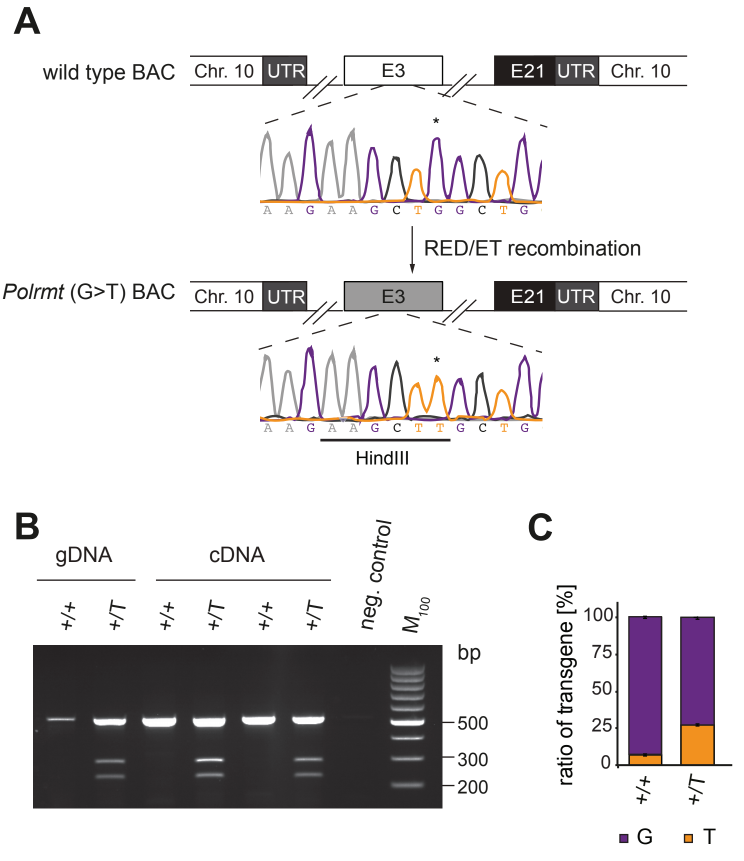


**Supplementary Figure 1 - Miranda et al**

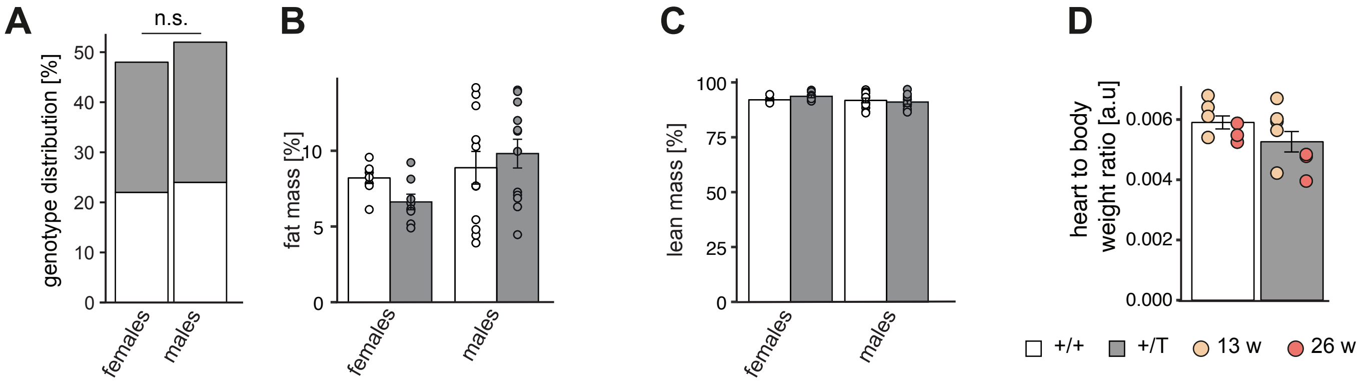


**Supplementary Figure 2 Miranda et al**

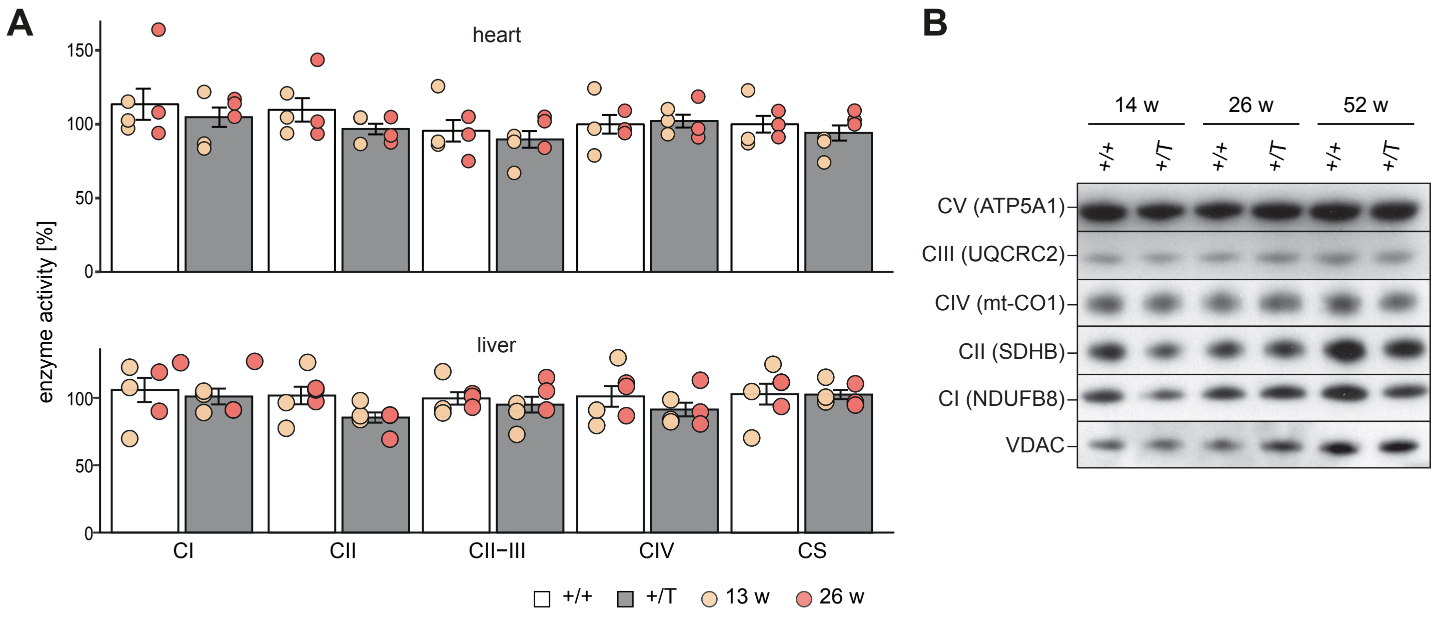


**Supplementary Figure 3- Miranda et al**

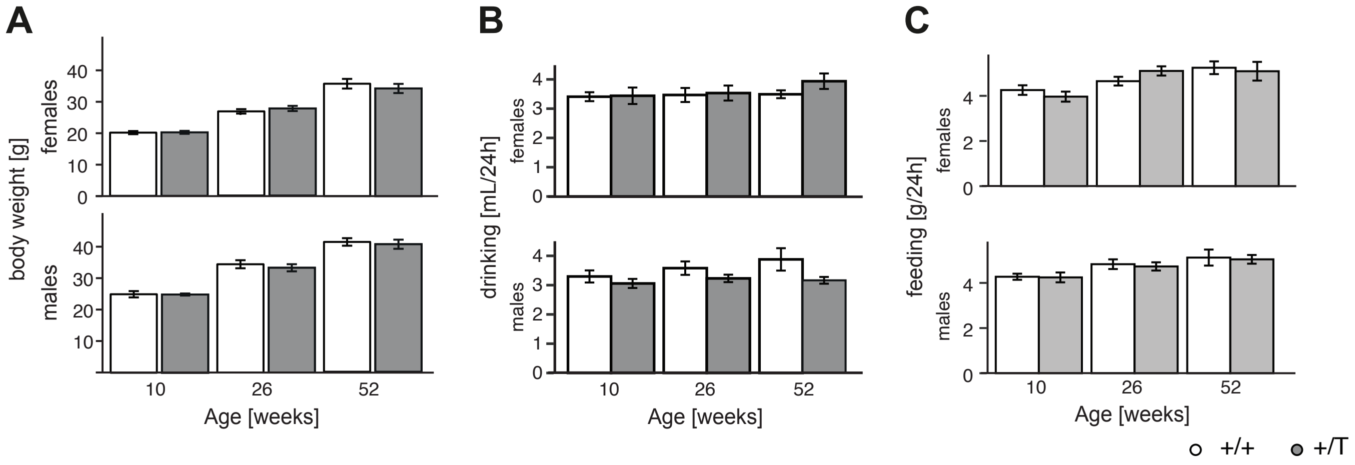


**Supplementary Figure 4- Miranda et al**
